# Can Living with an Alien Invasive Fish, *Tilapia*, Influence the Shoaling Decision-Making and Exploratory Behaviour of an Air-Breathing Freshwater Fish, the Climbing Perch?

**DOI:** 10.1101/839563

**Authors:** V V Binoy, Bhagyasree J Ingle, Aniket Bhattacharya, Anindya Sinha

## Abstract

The biodiversity of freshwater aquatic ecosystems is threatened by invasive alien species across the world. We studied the impact of the presence of an invasive piscine species, the tilapia *Oreochromis mossambicus* and acquisition of familiarity with it on the social decision-making and exploratory behaviour of a native, air-breathing, freshwater fish, the climbing perch *Anabas testudineus*. Our results reveal that the climbing perch did not show any significant preference or aversion to any of the stimulus shoals when unfamiliar monospecific shoals of tilapia, mixed-species shoals of tilapia and climbing perch that were divergent in the composition, or groups comprising only tilapia familiar to the subject fish for a duration of 30, 60, 90 or 120 days, were presented in opposition to a shoal with an equal number of unfamiliar conspecific individuals. No preference for isolated familiar individual tilapia was also observed against its unfamiliar counterpart or a conspecific individual. It is also noteworthy that the propensity of subject climbing perch to initiate exploration of a novel area (a measure of boldness) or exploratory activity and its sociability remained unchanged under different social conditions, including presence of unfamiliar conspecific, familiar conspecific, unfamiliar heterospecific or familiar heterospecific individuals. These results are discussed in the light of ever-increasing levels of invasion by alien fish species and the struggle for survival that currently confront native piscine species in most tropical freshwater ecosystems globally.

## Introduction

Intrusion of invasive alien piscine species into aquatic ecosystems and the subsequent alterations of the existing selection pressures on different levels of biological organisation (genetic, individual, population or community) remains one of the most important causes for worldwide extinction of indigenous fish fauna (Cucherousset and Olden 2011; Ellender and Weyl 2014; Gozlan 2015). Such extermination is often instigated either by the failure of indigenous piscine species to cope with the changes, caused by the invader, in the dynamics of the abiotic elements of the habitat or the inability of the local species to compete with the aliens in attaining biologically significant resources due to the lack of familiarity with the survival strategies followed by the latter (Cucherousset and Olden 2011; Gozlan 2015). Sometimes, attempts by the resident species to adapt to such novel situations by adjusting their own behaviours could negatively affect their survival and fitness (Blanchet *et al*. 2007, 2008; He *et al*. 2018). For instance, the presence of invasive bluegill *Lepomis macrochirus* and largemouth bass *Micropterus salmoides* was powerful enough to alter the normal behavioural patterns of the native Sacramento perch *Archoplites interruptus* and Cape galaxias *Galaxias zebratus*; the local inhabitants spent significantly more time under canopy cover, leading to the reduction in foraging and growth rates (Marchetti 1999; Shelton *et al*. 2008). Similarly, a recent report highlighted the modification of vital behaviours, such as aggregation, diurnal activity patterns and food preference of the native brown trout *Salmo trutta* in the presence of the invasive brook trout *Salvelinus fontinalis* (Larranaga *et al*. 2018).

If the invasive fish is social in nature, there is often a possibility that their interaction with the local shoaling species could lead to the emergence of mixed-species groups, particularly in habitats with limited space. According to Camacho-Cervantes *et al*. (2014, 2018), alien invaders gain foraging benefits by shoaling with native species and in many contexts, the latter also join the exotic fishes for social benefits. However, living with invasive species is not devoid of any cost; there are studies that indicate how the social life of indigenous species could be affected negatively by exotics (Blanchet *et al.* 2008). For instance, in the Atlantic salmon *Salmo salar*, the appearance of the exotic rainbow trout *Onchorynchus mykiss* led to interspecific competitive interactions that disrupted the dominance hierarchies and growth trajectories of the salmon, both at the individual and group level (Blanchet *et al.* 2008; Houde *et al.* 2015, 2016). Furthermore, such behavioural phenotypic changes appeared to be associated with the regulation of transcription of genes in the brain, including those implicated in protein turnover, neuronal structural change and oxygen transport (Roberge *et al.* 2008). Hence, the socialisation of indigenous and invasive fish species and the resulting formation of mixed shoals could pose a serious threat to many native species.

In fishes, the continuous exposure and interaction with conspecifics as well as with heterospecifics could lead to the development of familiarity (Paijmans *et al*. 2019). Many piscine species behave differently towards familiar and unfamiliar individuals and even show preference for shoals of familiar heterospecifics over groups of unfamiliar conspecifics (Ward *et al*. 2003; Coppocka *et al.* 2016; Kleinhappel *et al.* 2016). The impact of familiarity is not restricted to individuals; pairs of fish varying in their acquaintance with one another could also exhibit marked divergence in various behavioural patterns including exploratory behaviour and leadership (Lucon-Xiccato *et al*. 2019). Furthermore, dyads comprising of individuals from two different piscine species performed differently in situations requiring the utilisation of numerical ability and social decision-making (Bai *et al*. 2019). These results points to the need for studying the behaviour and cognitive abilities of mixed-species groups, especially those composed of both alien invasive and native species and varying in size and composition as well as the degree of familiarity between the constituent individuals. Such studies focusing on the behavioural adaptations of both native and invasive species (Seivers *et al*. 2012; Cote *et al*. 2013) can provide useful insights for the successful management of alien invasive fishes and the conservation of indigenous species in increasingly threatened aquatic ecosystems across the world.

Currently, one of the most important invasive piscine species anywhere in the world is the African mouth-brooder cichlid or the Mozambique tilapia *Oreochromis mossambicus* (Russell *et al.* 2012). According to Barki and Karplus (2016), this fish can significantly impact the behaviour of other aquatic animals present in their habitat. In India, the tilapia was introduced from Sri Lanka in 1952 and today, it is one of the most proliferating invasives in several major river systems across the subcontinent (Sugunan 2000; Lakra *et al*. 2008). In Kerala, the southernmost state of India, many natural water bodies, including those located in biodiversity hotspots (Raghavan *et al*. 2008), now harbour fast-growing populations of tilapia, who share these habitats with several native species, including the climbing perch *Anabas testudineus*. Our previous laboratory studies have demonstrated that climbing perch can acquire familiarity with a shoal of conspecific individuals and possess the ability to distinguish an isolated, familiar conspecific from an unfamiliar one; such familiarity subsequently influenced the shoaling decisions of the species (Binoy and Thomas 2006; Binoy *et al*. 2015). Hence, there is a strong likelihood that native populations of climbing perch could develop familiarity-dependent preference for shoals of invasive, alien heterospecifics and *vice versa* if the probability of them coming in contact is significantly high. Moreover, it also remains unknown what influences long-term interactions with invasive species could have on the behaviour and fitness of the climbing perch.

The current study explores the impact of the presence of the exotic tilapia and acquired familiarity with it on various aspects of climbing perch behaviour, including shoaling decisions and recognition of individual tilapia, as well as the behaviour of familiar and unfamiliar mixed-species dyads, comprising of climbing perch and tilapia, while exploring an open area. We also discuss the major implications of our findings for the conservation and management of the indigenous climbing perch in its freshwater range, increasingly being invaded by the tilapia, across the Indian subcontinent.

## Methods

### Study species and its husbandry

The climbing perch *Anabas testudineus* is an obligatorily air-breathing fish, inhabiting freshwater ecosystems of India and other southeast Asian countries (Talwar and Jhingran 1991). The study fish (6 ± 2.4 cm, mean ± SE, standard length, *L*_S_) were collected from the *Kole* paddy fields of Irinjalakuda (10.30°–10.42°N, 76.20°–76.28°E) in Thrissur district of the southern Indian state of Kerala and transferred to the laboratory. Tilapia was purchased from aquarium keepers and individuals of length 6.1 (± 2.8) cm were chosen for the experiments in this study. As some of our own earlier studies have indicated that familiarity with individual fish can influence shoaling decisions in this species (Binoy and Thomas 2006; Binoy *et al.* 2015), the absence of tilapia in the sites, from which the study climbing perch was collected, and in neighbouring areas was ensured by interviewing the local fishermen.

### Shoaling decisions: Apparatus and general procedures

Shoal-choice experiments were conducted in an apparatus consisting of an aquarium (85 × 32 × 32 cm), divided into two side chambers (16 × 32 × 32 cm) and a central chamber (53 × 32 × 32 cm) with transparent Plexiglas sheets with perforations (Binoy *et al.* 2015). Three sides of the aquarium were covered with black paper and water filled up to a height of 28 cm. The side chambers were used to present stimulus shoals while the subject fish was introduced individually in the central compartment in a transparent presentation cage (15 × 10 × 32 cm), made of acrylic sheets with perforations. Each subject fish was given six min to assess the stimulus shoals before it was released into the central chamber by raising the presentation cage. Time spent by the subject fish in the ‘preference zone’ (area within 5 cm from either of the stimulus shoals) during the test time (6 min) was taken as the indication of its preference for the shoal present in the adjacent chamber (Binoy *et al.* 2015). We used each subject fish only once.

### Experiment 1: Influence of shoal composition on social decision-making by climbing perch

These experiments examined the shoal-choice behaviour of subject fish when provided with mixed-species shoals comprising of unfamiliar conspecific individuals and tilapia that varied in species composition in opposition to an equal-sized monospecific shoal of climbing perch. The different combinations of the four-individual mixed-species shoal tested were: four climbing perch, one tilapia + three climbing perch, two tilapia + two climbing perch or three tilapia + one climbing perch. The second stimulus shoal in all these experiments consisted of 4 unfamiliar climbing perch. The shoaling decision-making by the climbing perch was also tested by providing four unfamiliar tilapia against four unfamiliar climbing perch as the two simultaneous stimuli. The experimental procedure was exactly as described in the section on Shoaling decisions: Apparatus and general procedures, given above. We tested 20 individuals in each of these experiments.

### Experiment 2: Time required for climbing perch to acquire familiarity-dependent preference for a shoal of tilapia

A total of 12 fish—six climbing perch and six tilapia—were allocated to each of the familiarisation aquaria (80 × 40 × 45 cm). The water level of these aquaria was kept at a height of 40 cm. In order to avoid the confounding effect of the body size of shoal-mates on their shoaling decisions (Griffiths and Ward 2011), only size-matched individuals were used in these experiments. Each aquarium was covered with black paper on three sides and steel grids placed on the top to prevent the fish from jumping out of the water. Water temperature was maintained at 25 (± 1)°C and light hours at 12L:12D. The fish were fed twice (morning and evening) and the excess food pellets siphoned out 30 min after the start of each feeding session (Binoy *et al*. 2015).

Different groups of climbing perch were allowed to familiarise with shoals of heterospecific individuals for durations of 30, 60, 90 or 120 days respectively. The impact of acquired familiarity on the shoaling decision of the subject climbing perch was tested by presenting a shoal of five unfamiliar conspecific individuals against an equal-sized shoal of tilapia, varying in the durations of familiarisation with the subject fish of 30, 60, 90 or 120 days. Eighteen individual climbing perch were tested in each of these shoal-choice experiments.

### Experiment 3: Can acquired familiarity help climbing perch to recognise individual member of heterospecific shoals?

The ability of climbing perch to recognise an individual heterospecific, based on prior experience, was also tested by presenting single stimulus fish in each of the side chambers of the testing aquaria. The different combinations of the stimuli fish tested included unfamiliar conspecific vs. unfamiliar heterospecific, unfamiliar conspecific vs. familiar heterospecific, unfamiliar heterospecific vs. unfamiliar heterospecific or unfamiliar heterospecific vs. familiar heterospecific. In the experiments with familiar heterospecifics, each of the subject fish was familiarised with the respective stimulus tilapia for 120 days. We tested 16 individual climbing perch in each of these experiments.

### Experiment 4: Decision-switching while choosing shoals

It is now well established that individual fish switch between the stimulus groups available while preforming their shoal choice. According to Lucon-Xiccato *et al*. (2017a) such shuttling between the stimulus shoals helps the fish to assess the shoals available before taking an appropriate decision. Bai *et al*. (2019) have associated the shuttling behaviour of the individual with its personality trait of boldness while in the opinion of Rogers *et al*. (2011), this behaviour is highly correlated with the body size, learning ability and predation pressure experienced by the fish. We quantified the number of shuttles exhibited by the subject fish between the preference area near one stimulus shoal to the other in all experiments included under the Experiments 1, 2 and 3 described above.

### Experiment 5: Influence of the presence of familiar and unfamiliar conspecific and heterospecific individuals on specific personality traits of climbing perch

Social piscine species are now known to behave differently in novel situations if accompanied by familiar or unfamiliar individuals (Lucon-Xiccato *et al*. 2017b). These studies mainly focused on the behaviour of monospecific pairs or groups in novel environments (Lucon-Xiccato *et al*. 2019). Similar studies exploring the behaviour of mixed-species dyads, comprising individuals from both indigenous and invasive species and the impact of the acquisition of familiarity between them are, however, scanty in the literature although the potential of such investigations in contributing to the development of effective strategies to control invasive piscine species is enormous.

The influence of a conspecific or invasive heterospecific companion on the exploratory behaviour and sociability of the climbing perch was measured using an apparatus consisting of an open area connected with a refuge via a removable partition (Lucon-Xiccato *et al*. 2017b). The apparatus that we used in our study was made up of an aquarium (70 × 32 × 32 cm) divided into two compartments—Chamber A (20 × 32 × 32 cm) and Chamber B (50 × 32 × 32 cm)—by an opaque plexiglass sheet with a guillotine door (10 × 7 cm; length × height) in the centre (Binoy *et al.* 2015). Three sides of the aquarium were covered using black paper while Chamber A, which acted as both start chamber, and the refuge was made opaque by keeping black acrylic sheet on the open sides. However, the top of the start chamber was not covered, as it could affect the exploratory behaviour of the climbing perch. The apparatus was illuminated using a compact florescent lamp (32 W), suspended above the assembly.

The behaviour of the isolated climbing perch was tested as follows: each subject fish was introduced into Chamber A individually and the latency to initiate exploration—defined as the time taken by the focal fish to come out of the start chamber through the door provided—of Chamber B was recorded. The subject fish that moved to Chamber B were given six min each for exploration and the time spent stationary there (measure of activity) was documented. Another behaviour quantified was the number of air-gulps during the exploration of Chamber B. Those individuals that failed to emerge from Chamber A after six min were allocated a ceiling value of 360 seconds and the trial then terminated. This protocol was exactly repeated in another set of experiments, in which the focal climbing perch was introduced into the start chamber together with an unfamiliar conspecific, familiar conspecific, unfamiliar heterospecific tilapia or a familiar tilapia. Along with the earlier measures of boldness and activity, the time spent by the focal fish within an area of two cm around the stimulus fish—a measure of sociability / shoal cohesion (Lucon-Xiccato *et al*. 2017b)—was also estimated in these sets of experiments. We tested 25 different individual climbing perch in each trial with the individual fish being used in one experiment never employed in any other experiment.

## Analysis

The Preference Index for each test fish used in Experiments 1, 2 and 3 was calculated as the proportion of time spent by each subject near the conspecific shoal or individual divided by the total time spent near each stimulus shoal or individual (Gómez-Laplaza and Gerlai 2016). One-way ANOVA was used to test the influence of various social situations on shoal choice by the test fish in each of the Experiments 1, 2 and 3 as well as to compare the decision-switching in these experiments as these data were established to be normally distributed. Tukey’s test was utilised for the *post hoc* analysis of the tests in Experiment 2.

The data obtained from Experiment 5 did not follow a normal distribution even after transformation. Hence, the non-parametric Kruskal-Wallis test was used to examine the influence of the presence of familiar and unfamiliar conspecific and heterospecific individual on the boldness, activity, shoal cohesion and air-gulping of the subject fish in these experiments.

## Results

Our study revealed that climbing perch failed to exhibit a significant preference to any of the stimuli shoals, composed of four conspecific individuals, four heterospecific tilapia, three tilapia + one climbing perch, two tilapia + two climbing perch or one tilapia + three climbing perch against a group of four climbing perch (ANOVA, F_4,100_ = 1.11; p = 0.11; Fig. 1). Exposure for 30, 60, 90 or 120 days also did not help the subject fish to develop a familiarity-dependent preference for a shoal of tilapia over an equal-sized group of unfamiliar conspecific individuals (F_4,85_ = 0.71, p = 0.59). Moreover, no preference was evident when the choices given to the test fish were isolated individual conspecific vs. unfamiliar heterospecific, unfamiliar conspecific vs. familiar heterospecific or unfamiliar heterospecific vs. familiar heterospecific individuals (F_3,64_ = 0.75, p = 0.52; Fig. 2). Akin to the previous results, no significant variation was also observed in decision-switching by the test fish—the number of times the subject fish changed its preference by moving between the stimuli available in the side chambers during testing time—in the diverse social contexts provided in Experiments 1, 2 and 3 (F_13,262_ = 1.28, p = 0.22).

**Figure 1.**
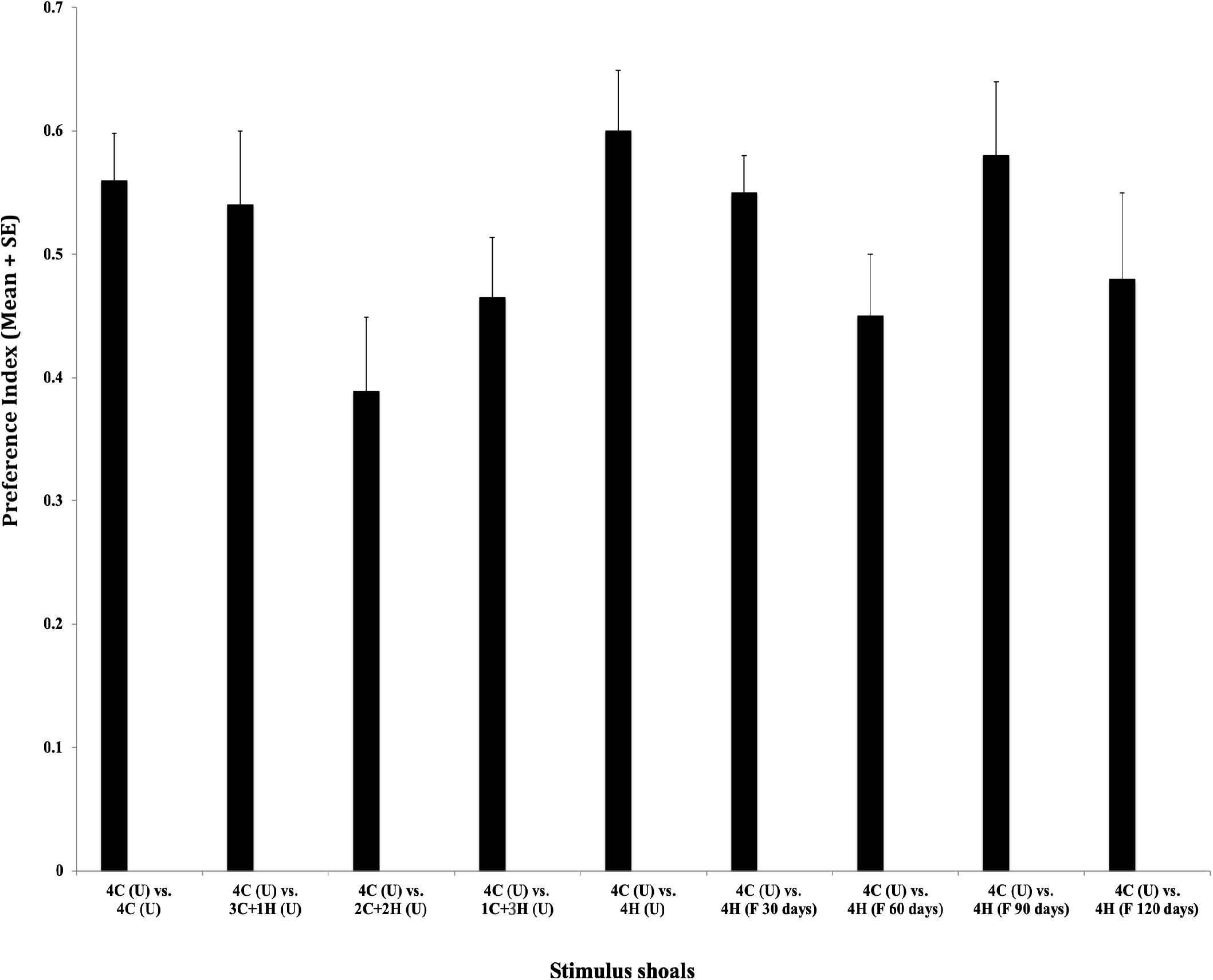
Influence of species composition of the stimulus shoals and individual experience with shoals of heterospecific tilapia over varying periods of time on shoaling decisions of climbing perch. The number of conspecific climbing perch individuals (C) and heterospecific tilapia individuals (H) in the two stimulus shoals and the duration of familiarity with monospecific shoals of heterospecific tilapia (F) that the subject fish had to choose between has been indicated below each bar. The letter U denotes unfamiliarity with the stimulus shoals in all cases. The Preference Index has been defined in the text.

**Figure 2.**
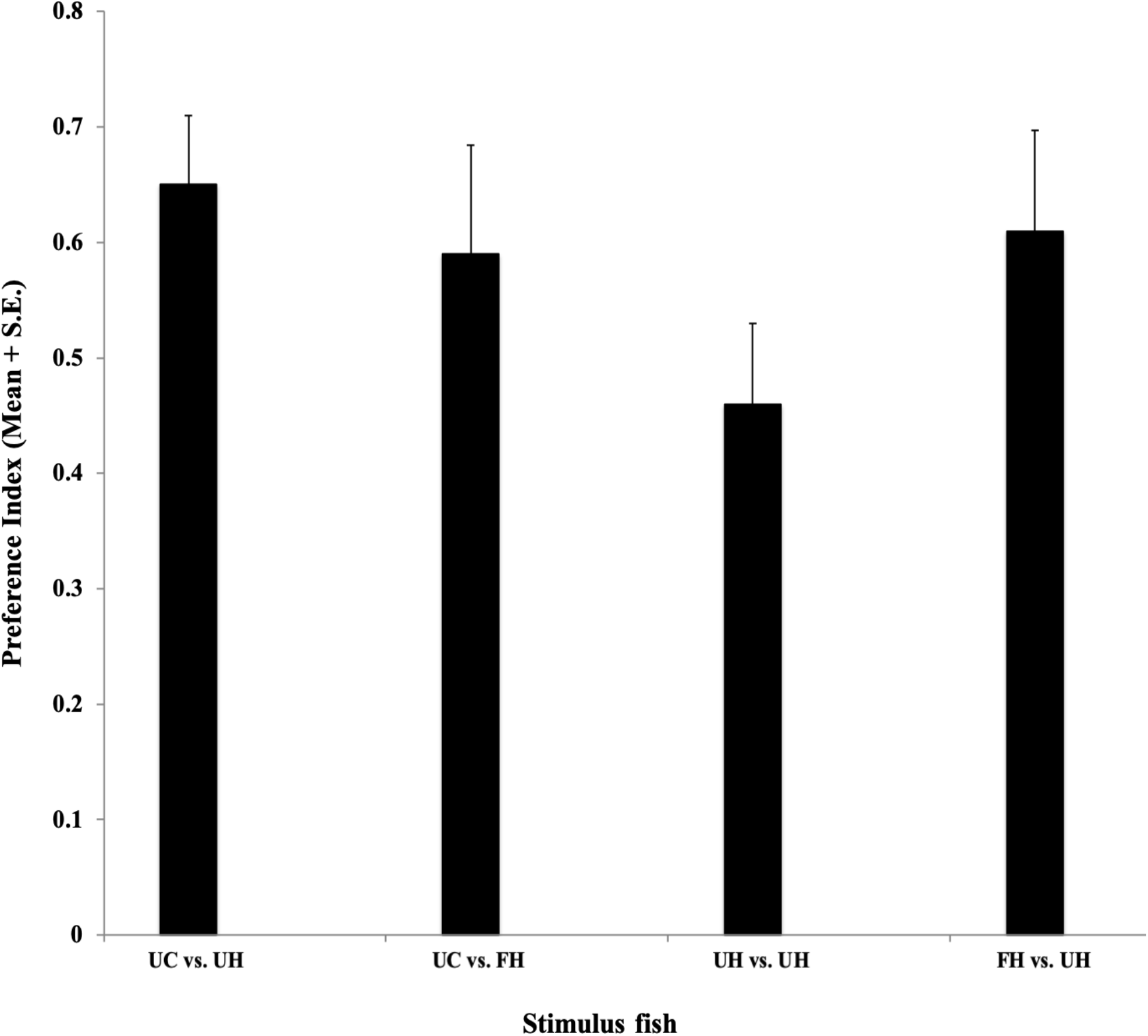
Influence of acquired familiarity with isolated conspecific climbing perch and heterospecific tilapia individuals on shoaling decisions of climbing perch. UC: Unfamiliar conspecific individual; UH: Unfamiliar heterospecific individual; FH: Familiar heterospecific individual. The Preference Index has been defined in the text.

**Figure 3.**
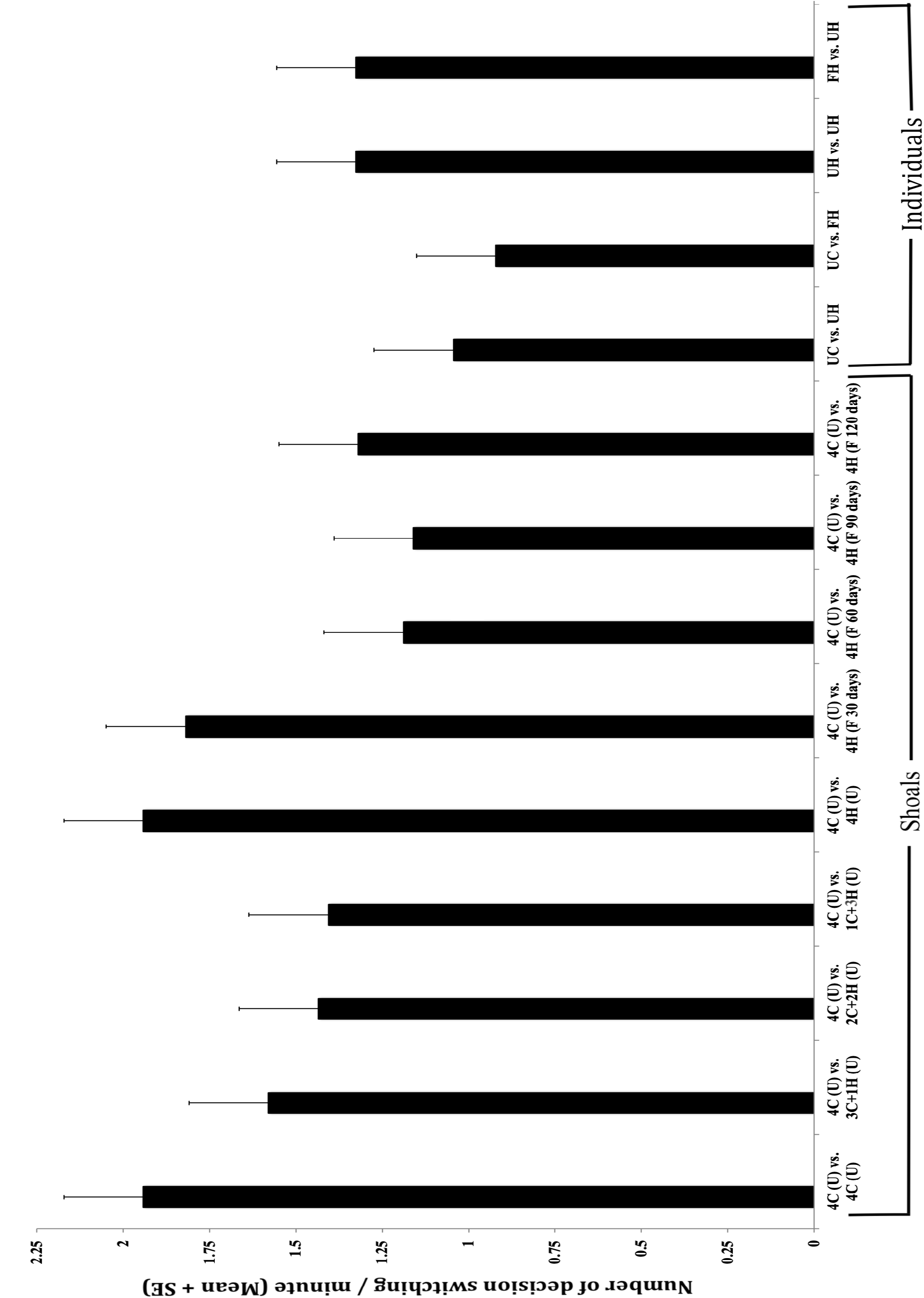
Frequency of decision-switching — the number of times the subject fish changed its preference by alternately moving between the stimuli available in the side chambers per minute of testing time — exhibited by the climbing perch in the different social contexts tested. The number of conspecific climbing perch individuals (C) and heterospecific tilapia individuals (H) in the two stimulus shoals and the duration of familiarity with monospecific shoals of heterospecific tilapia (F) that the subject fish had to choose between has been indicated below each bar. The letter U denotes unfamiliarity with the stimulus shoals in all cases. UC: Unfamiliar conspecific individual; UH: Unfamiliar heterospecific individual; FH: Familiar heterospecific individual.

The subject climbing perch did not display any statistically significant difference in the time taken to leave the start chamber and initiate explorations of Chamber B, an apparent measure of boldness, either in isolation or in the presence of conspecific or heterospecific individuals, unfamiliar or familiar (Kruskal-Wallis test, χ^2^ = 8.30, n = 25, p > 0.05; Fig. 4A). Moreover, analogous to boldness, there was no significant variation, across different social contexts tested, in the time spent in exploration of the swim-way (χ^2^ = 8.93, n = 25, p > 0.05 Fig. 4B), in sociability towards stimulus fish (χ^2^ = 3.78, n = 25, p > 0.05; Fig. 4C) or in the frequency of air-gulps (χ^2^ = 2.16, n = 25, p > 0.05; Fig. 4D) exhibited by the test climbing perch individuals.

**Figure 4.**
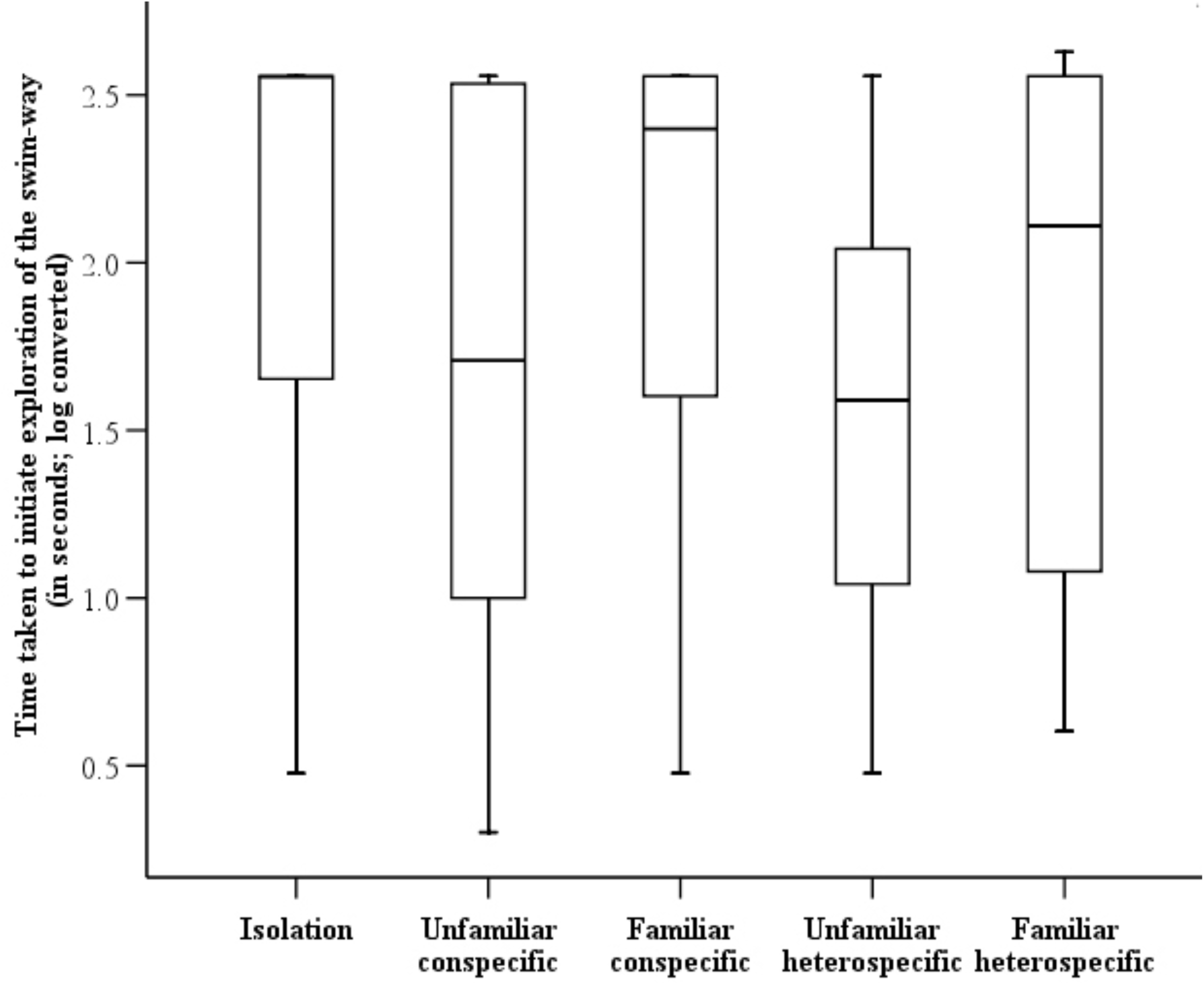
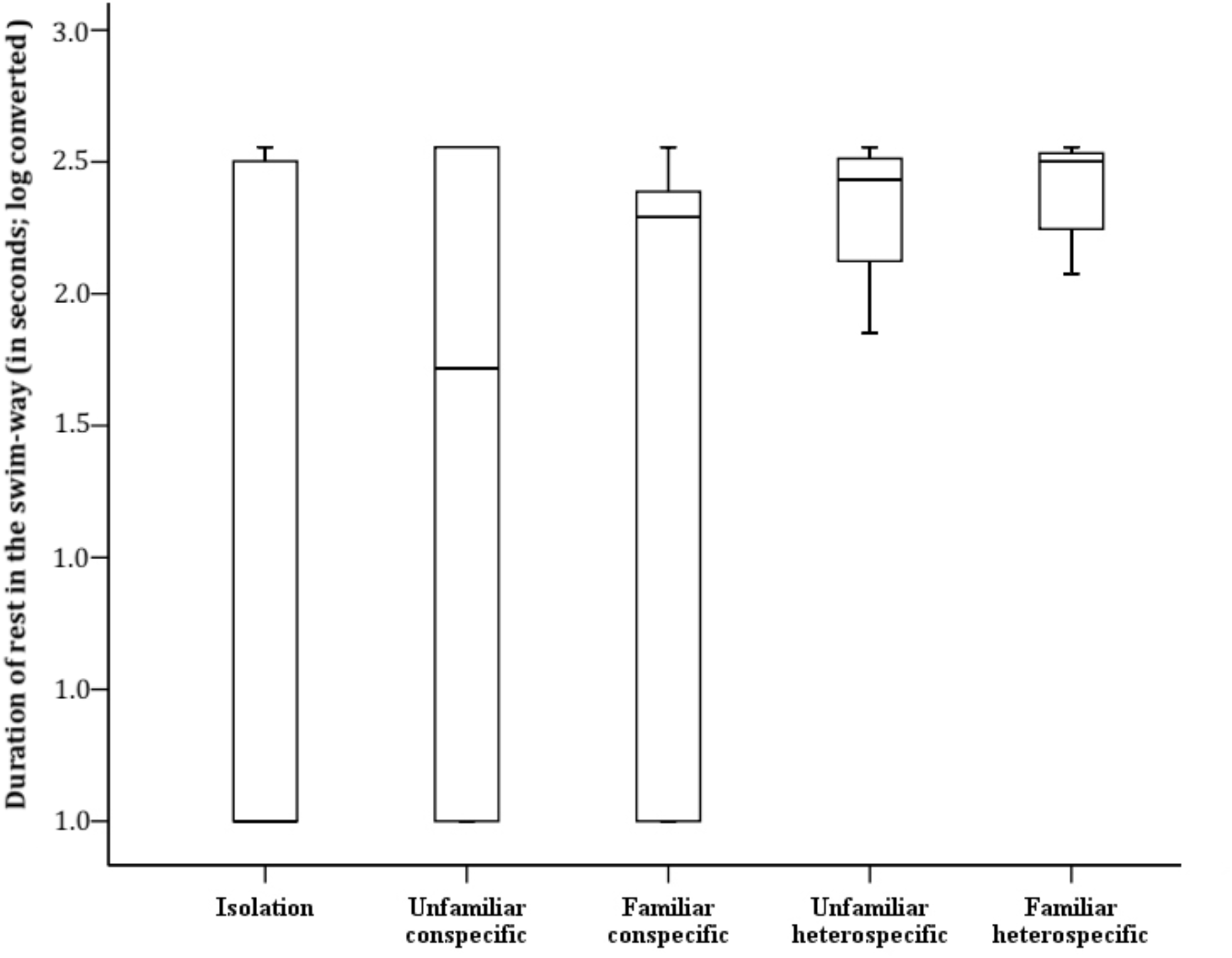
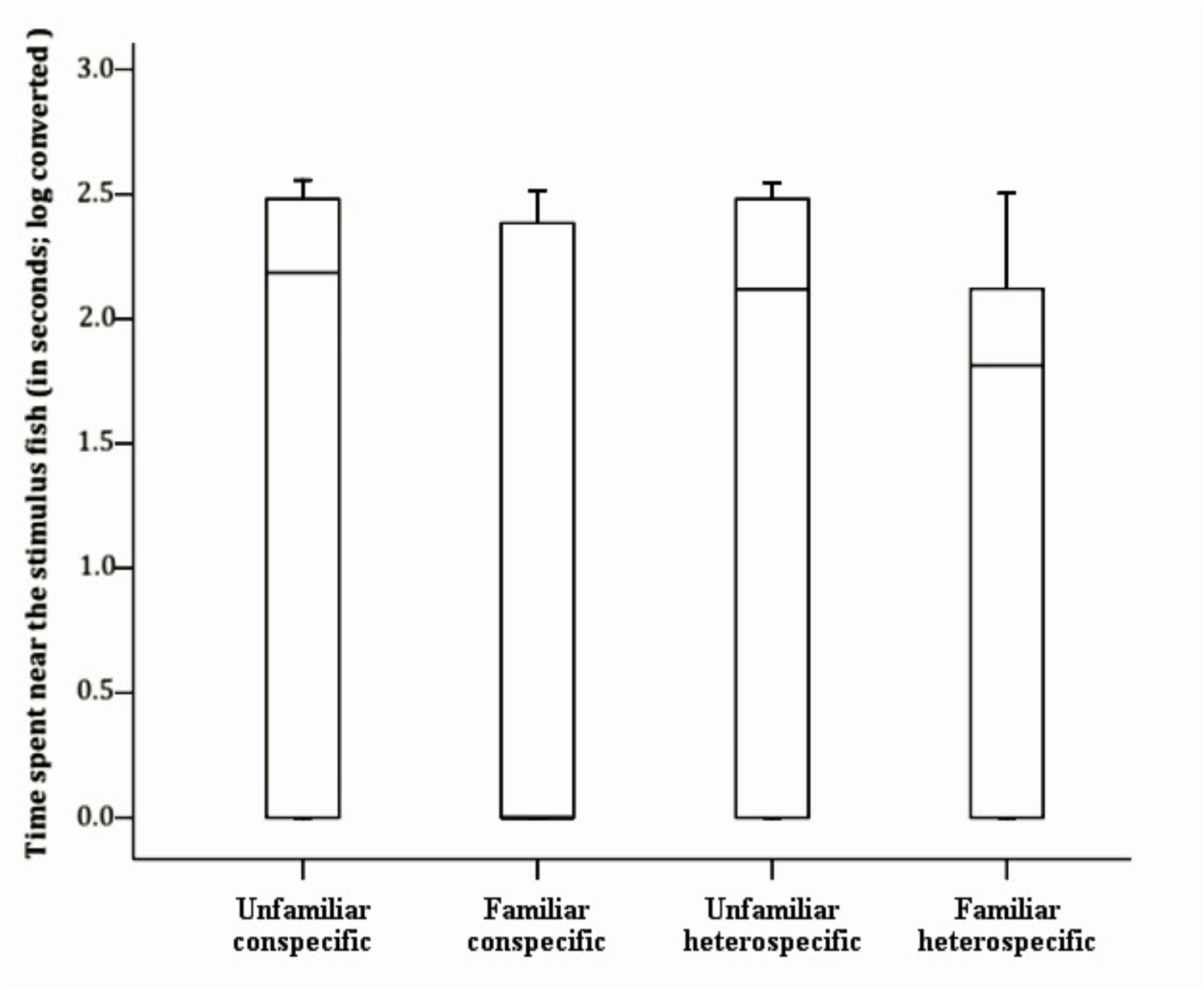
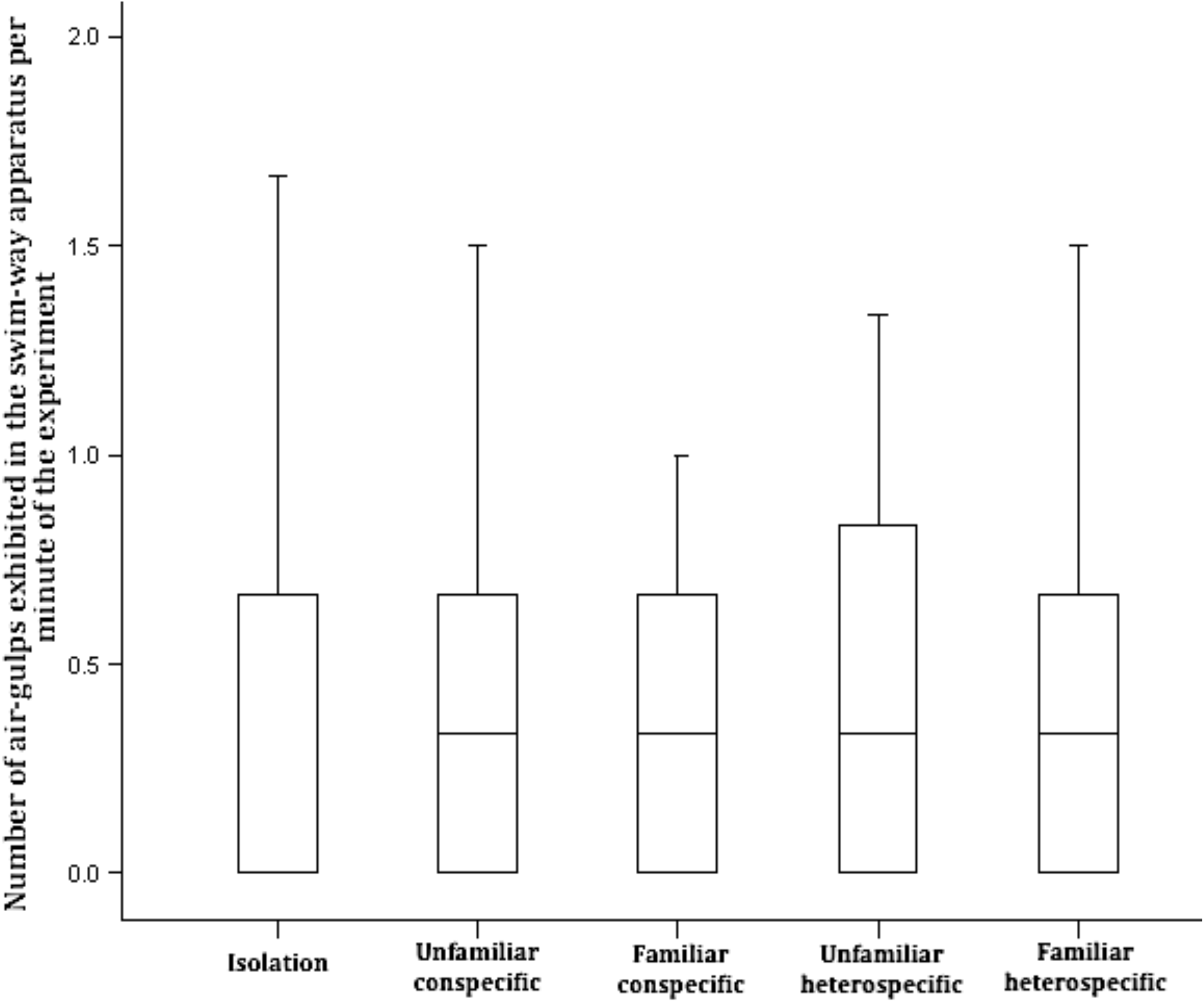
Measures of boldness (A), behavioural activity (B), sociability (C) and air-gulping (D) displayed by climbing perch, either in isolation or in the presence of unfamiliar or familiar conspecific or heterospecific individuals. Boldness was measured as the time taken to initiate exploration of the swim-way, behavioural activity as the time spent in rest while exploring the swim-way, sociability as the time spent within 2 cm of the stimulus fish, and air-gulping as frequency of air-gulps per unit time spent in exploration. Note that sociability, by definition, could not be measured in isolation. The data have been represented as median quartiles.

## Discussion

Social animals are believed to typically take decisions to leave or join conspecific groups as a resultant of the costs to be borne in terms of competition for biologically significant resources, attraction of the group to predators or chances of acquiring infection in the group as against the foraging or anti-predator benefits obtained by being a member of such aggregations (Ward and Webster 2016). More proximately, various group-living piscine species appear to be influenced by different factors, ranging from the body size of shoal-mates to their competitive abilities, when they take social decisions (Griffiths and Ward 2011). Although joining a shoal of conspecific individuals could be potentially advantageous due to their shared morphology and behaviour, which could translate into efficient group manoeuvres or other antipredator benefits, many fishes are often seen to form mixed-species groups (Quartini *et al*. 2018; Smith *et al*. 2018; Paijmans *et al*. 2019), including those with alien invasive species (Camacho-Cervantes *et al*. 2018). While living with heterospecifics who are divergent in their body shape or colour or in specific behavioural traits, can still be advantageous in terms of the dilution effect against predation, membership of such groups can also be associated with the costs of being an ‘odd’ (Landeau and Terborgh 1986). Individuals that appear or behave differently in a group could attract the unwanted attention of predators and thus put themselves or other members of the group in danger (Rogers *et al*. 2011; Quattrini *et al*. 2018). In the current study, our subject species, the climbing perch, did not show any significant variation in its social preference when presented with a choice of equal-sized shoals of unfamiliar conspecifics and heterospecific tilapia. Although several studies have shown that fishes often do exhibit a reduced preference for shoals containing odd individuals (Quattrini *et al*. 2018), our subject fish failed to display any preference or aversion towards the mixed-species shoals, comprising climbing perch and tilapia, but varying in the species composition, which they were tested with.

We had earlier noted that a familiarity-based preference for a shoal of conspecific individuals was acquired by climbing perch after 14 days of familiarisation (Binoy and Thomas 2004; Binoy *et al.* 2015) while the species was also capable of distinguishing isolated conspecific individuals of a familiar shoal from those of unfamiliar ones (Binoy and Thomas 2006). These, we believe, are abilities that may have evolved to be essential in avoiding unwanted aggression and, at the same time, enhance the potential benefits of shoal life. Living with tilapia for long periods of even 120 days, however, neither helped the climbing perch to develop a preference for heterospecific individuals nor did the subject fish exhibit a bias towards familiar heterospecific over unfamiliar heterospecific or unfamiliar conspecific individuals. Moreover, the lack of significant variation in the frequency of decision-switching—”the frequent movement that fish display across the available stimulus shoals in order to sample them and make an ultimately beneficial choice” (Jones *et al*. 2010)—shown by our climbing perch in the various social contexts tested may be taken as an indication of the ambivalence displayed by the subject fish towards conspecific and exotic heterospecific individuals. According to Quattrini *et al*. (2018), individual fish may take a decision to join a shoal of heterospecifics if it experiences relatively fewer phenotypic differences between its own and the other species. Therefore, if the climbing perch perceives a tilapia as a fish totally different from its conspecifics, there is clearly a good chance that the neophobia stimulated by the heterospecific fish could induce the subject individuals not to sample the stimulus shoals available in the side chambers of the test apparatus (see also Bai *et al*. 2019).

The lack of a species-specific shoaling preference displayed by the climbing perch, its not necessarily being aversive to mixed-species groups varying in the number of conspecific or alien invasive heterospecific individuals or its inability to develop familiarity-dependent preference towards groups of or towards individual tilapia raise many ecological concerns. These traits may indicate the possibility that climbing perch individuals may leave their own shoals and join group of tilapia, when shoals of these species encounter one another in their natural habitats. Such a decision would, of course, lead to the newly-formed mixed-species shoals to face foraging challenges of different kinds and levels of predation pressures in these waterbodies. The climbing perch is an obligatory air-breathing fish and has to surface frequently to gulp atmospheric air, a behaviour that makes it rather vulnerable to aerial predation (Kramer *et al*. 1983; Norris 1994). Air-breathing fish typically reduce this indispensable risk of predation by orchestrating their surfacing with shoal-mates, thus potentially confusing the aerial predator, which finds it difficult to eye-lock with a single individual amidst a rapidly moving group, in order to initiate the attack (Sloman *et al*. 2009). There is thus a good possibility that surfacing to breathe alone or in a relatively small group could become a costly exercise for the climbing perch, when shoaling with the non-air-breathing tilapia.

According to Galhardo *et al.* (2012), males of tilapia *Oreochromis mossambicus* exhibit more exploratory behaviour and less neophobia in the presence of a familiar rather than an unfamiliar conspecific. However, the company of tilapia, familiar or unfamiliar, did not appear to induce any marked variation in the hesitation to initiate exploration of a novel environment, time spent in exploring it or in the frequency of air-gulping by climbing perch. A similar response was exhibited by the species when it was tested for these behavioural traits in the presence of known or unknown individuals of its own species. The consistent sociability displayed by the climbing perch towards both conspecific and heterospecific individuals, before and after familiarisation with them over relatively long durations of time, however, also supports our hypothesis that isolated members of this species may readily join heterospecific individuals, including alien invasive heterospecific tilapia, if they have enough opportunities to encounter them.

The results obtained in our current laboratory-based study, nevertheless, need to be validated in natural habitats, as species-specific assortment, active avoidance of odd individuals through passive or active exclusion as well as a marked preference to join a group comprising conspecific individuals, could become strong when the presence of predators is sensed by the members of a shoal (Wolf 1985; Griffiths and Ward 2011). Hence, more extensive investigations of the decisions made by climbing perch with regard to their shoaling or managing oddity in mixed-species shoals, if indeed they form in natural environments, especially in the face of foraging challenges or the presence of predators, are essential in order to address questions on potentially adaptive behavioural strategies displayed by the species to ensure its fitness in habitats facing extensive invasion by exotic piscine species.

To conclude, it is entirely conceivable that patterns of behavioural interaction between invading, alien species and resident fish species can change over time as they acquire social experience with one another. In this context, it is important to note that individual climbing perch migrate from one water body to another during the monsoon rains (Sokeng *et al.* 1999) and being a social species, individuals reaching unfamiliar waterbodies would require to find shoal-mates and reside with them. Populations of tilapia are dramatically increasing in natural inland waterbodies across the Indian subcontinent due to the introduction of this economically important species by the government and private agencies to enhance revenue from the fisheries sector (Roshni *et al*. 2016). We, however, still do not have adequate information on the patterns of interaction between these species in natural habitats or on the selection pressures and fitness advantages that mixed-species grouping could impose on these two fishes. Hence, long-term studies focussing on the strategies utilised by the respective species to reduce the oddity effect, novel adaptations that may have evolved to bring in effective anti-predator strategies under different circumstances or the possible modifications of different species-typical social behaviours, displayed by both tilapia and climbing perch in both natural environments and under controlled laboratory conditions, are absolutely imperative in the near future. Such investigations would not only allow for a comprehensive understanding of the mechanisms of invasion by alien piscine species and their impacts on the native fish taxa (Juette *et al*. 2014; Berthon 2015) but also aid in the design and implementation of effective strategies to control the spread of the alien species in new habitats while managing the often rapidly affected and declining populations of important and rare indigenous species.

## Acknowledgements

VVB is grateful to the Science and Engineering Research Board (SERB), Department of Science and Technology, Government of India, for a Grant (SB/FT/LS-155/2012) that enabled this study. The experiments reported in this paper comply with the current relevant laws of India.

